# Exercise preserves fitness capacity during aging through AMPK and mitochondrial dynamics

**DOI:** 10.1101/2022.06.20.496837

**Authors:** Juliane Cruz Campos, Luiz Henrique Marchesi Bozi, Annika Traa, Alexander M van der Bliek, Jeremy M. Van Raamsdonk, T. Keith Blackwell, Julio Cesar Batista Ferreira

## Abstract

Exercise is a nonpharmacological intervention that improves health during aging, and a valuable tool in the diagnostics of aging-related diseases. In muscle, exercise transiently alters mitochondrial functionality and metabolism. Mitochondrial fission and fusion are critical effectors of mitochondrial plasticity, which allows a fine-tuned regulation of organelle connectiveness, size and function. Here we have investigated the role of mitochondrial dynamics during exercise in the genetically tractable model *Caenorhabditis elegans*. We show that in body wall muscle a single exercise session induces a cycle of mitochondrial fragmentation followed by fusion after a recovery period, and that daily exercise sessions delay the mitochondrial fragmentation and fitness capacity decline that occur with aging. The plasticity of this mitochondrial dynamics cycle is essential for fitness capacity and its enhancement by exercise training. Surprisingly, among longevity-promoting mechanisms we analyzed, constitutive activation of AMPK uniquely preserves fitness capacity during aging. As with exercise training, this benefit of AMPK is abolished by impairment of mitochondrial fission or fusion. AMPK is also required for fitness capacity to be enhanced by exercise, with our findings together suggesting that exercise enhances muscle function through AMPK regulation of mitochondrial dynamics. Our results indicate that mitochondrial connectivity and the mitochondrial dynamics cycle are essential for maintaining fitness capacity and exercise responsiveness during aging, and suggest that AMPK activation may recapitulate some exercise benefits. Targeting mechanisms to optimize mitochondrial fission and fusion balance, as well as AMPK activation, may represent promising strategies for promoting muscle function during aging.

**Significance Statement:** Exercise is a powerful anti-aging intervention. In muscle exercise remodels mitochondrial metabolism and connectiveness, but the role of mitochondrial dynamics in exercise responsiveness has remained poorly understood. Working in *Caenorhabditis elegans*, we find that the mitochondrial dynamics cycle of fission and fusion is critical for fitness capacity, that exercise delays an aging-associated decline in mitochondrial connectiveness and fitness capacity, and that the mitochondrial dynamics cycle is required for the latter benefit. AMPK, which regulates mitochondrial dynamics, is needed for exercise to maintain fitness capacity with age and can recapitulate this exercise benefit. Our data identify the mitochondrial dynamics cycle as an essential mediator of exercise responsiveness, and an entry point for interventions to maintain muscle function during aging.

## Introduction

Age-related disorders are a major and growing public health problem, and an epidemic of aging-related diseases has followed increases in both unhealthy living habits and life expectancy (1). This has made it critical to develop interventions that promote healthy aging, including improvements in healthspan – the period of life in which an individual maintains good health.

Impairment of mitochondrial quality control and the consequent accumulation of fragmented and dysfunctional mitochondria are pivotal in the establishment and progression of chronic degenerative diseases (2-4). Mitochondrial fission and fusion, commonly referred as mitochondrial dynamics, are essential for maintaining mitochondrial function. While fission segregates mitochondria into spherical organelles (5, 6), fusion allows mitochondrial components to be reconstituted within a new mitochondrial network (7), thereby facilitating mitochondrial metabolic remodeling and quality control. By functioning in a continuous cycle, these processes allow mitochondrial physiology to be maintained and fine-tuned (8). Indeed, genetic disruption of mitochondrial dynamics that impairs mitochondrial functions ultimately results in age-related disorders (9-12).

Exercise has been widely employed to improve quality of life and protect against degenerative diseases. In humans a long-term exercise regimen reduces overall mortality (13), and fitness capacity can be a valuable parameter for diagnostics of age-related diseases that include sarcopenia, osteoporosis, and cardiovascular and neurodegenerative diseases (14). Fitness capacity is also a valuable physiological marker of healthy aging (15), and its impairment is associated with poor disease prognosis and reductions in quality of life and survival across species (16-18). The various benefits of exercise are conferred in part through transient increases in energy expenditure that affect mitochondrial metabolism and network morphology (19). In rodents exercise enhances cardiac function in heart failure by favoring mitochondrial fusion (20). Disruption of mitochondrial fusion reduces exercise performance (23). Similarly, mitochondrial fission is necessary to increase bioenergetic flux and meet the cardiac and skeletal muscle energetic demand of exercise (21, 22). However, the contribution of integrated mitochondrial dynamics to fitness capacity, exercise responsiveness, and maintenance of fitness capacity during aging remains largely to be determined. The dynamic nature of mitochondria may have contributed to this gap of knowledge, because addressing these questions requires an experimental model in which these questions can be explored in real time in the setting of exercise, and over the lifespan of an organism.

The nematode *Caenorhabditis elegans* (*C. elegans*) is a powerful model for aging research because of its short lifespan, and amenability to genetic analysis and microscopy of living tissue (24). Notably, *C. elegans* also exhibits key features of the mammalian response to exercise (25-30). In these animals a single session of swimming leads to increased oxygen consumption, fatigue, and transcriptional changes towards mitochondrial oxidative metabolism (28), and daily exercise sessions improve health parameters across multiple tissues and can increase stress resistance and lifespan (25-27, 29, 30). Together with similar findings in *D. melanogaster* (31, 32), these studies suggest that fundamental exercise adaptations are conserved from invertebrates to humans. Here, we investigated the role of mitochondrial dynamics in fitness capacity and the benefits of exercise in young and aging *C. elegans*. We also explored how exercise capacity and benefits are affected by established anti-aging interventions, including activation of the energy sensor AMPK (AMP-activated protein kinase).

## Results

### Effects of aging and exercise on mitochondrial connectivity and fitness capacity

We first investigated how aging affects fitness capacity and mitochondrial dynamics in wild type (WT) worms. When we examined fitness capacity by recording animals during a swimming session, we observed a progressive decline in the number of body bends per second at 5, 10 and 15 days of adulthood compared to day 1, demonstrating that fitness capacity declines during aging (Fig. 1*A*). We also scored mitochondrial connectiveness in body wall muscle, which exhibits many similarities to mammalian skeletal muscle (28), based upon five classes that reflect a gradual increase in fragmentation and disorganization (Fig. 1*B*). At day 1 most muscle cells (76%) exhibited connected mitochondria (class 1), while at days 5 and 10 there was a significant and progressive shift towards fragmented/disorganized mitochondria (classes 3-5) (Fig. 1*B*). These results align with previous studies showing reduced fitness capacity and increased mitochondrial fragmentation during aging across species (17, 33-36).

**Figure 1.**
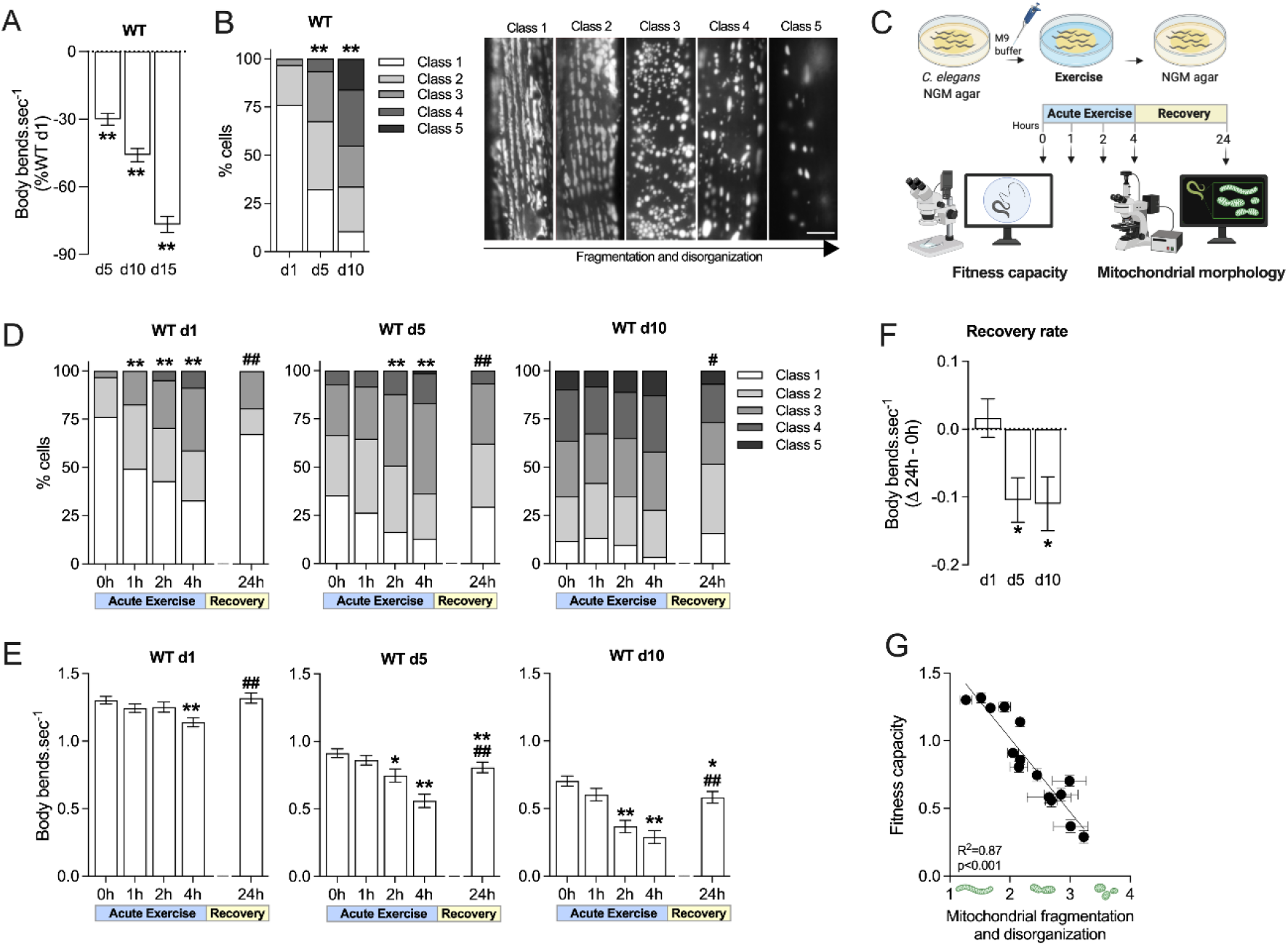
Effects of exercise and aging on mitochondrial dynamics. (A) Fitness capacity decay and (B) mitochondrial morphology in body wall muscle cells of WT worms during aging, with representative images of the 5 classes of mitochondrial network organization shown (similar to scoring in ref (29), Scale bar, 5 μM). (C) Acute exercise protocol: worms maintained at 20°C were subjected to 4 hours of swimming followed by 24 hours of recovery on agar plates. (D) Mitochondrial morphology in body wall muscle cells, (E) fitness capacity and (F) recovery rate of WT worms that were subjected to acute exercise on days 1, 5 and 10 of adulthood. Recovery rate refers to the difference in body bends sec^−1^ between 24h and the start of exercise (0h). (G) Correlation between fitness capacity and mitochondrial morphology of WT worms subjected to acute exercise, including data from d1, d5 and d10 of adulthood. Data are presented as mean ± SEM. *p <0.05 and **p <0.001 vs. WT (d1 or 0h); #p <0.05 and ##p <0.001 vs. 4h. Detailed statistical analyses, number of biological replicates and sample size are described in *SI Appendix*, Table S1.

Considering that exercise immediately triggers muscle energy imbalance (37), a major regulator of mitochondrial dynamics (38, 39), we investigated how the mitochondrial network is affected by a single cycle of exercise and recovery (Fig. 1*C*). Acute exercise induced a progressive increase in muscle mitochondrial fragmentation in young adult worms (day 1, Fig. 1*D*) along with a decrease in fitness capacity at the last hour of exercise (4h, Fig. 1*E*). Importantly, a 24-hour recovery period was sufficient to reestablish mitochondrial connectiveness (towards tubular and interconnected) (Fig. 1*D*) and mitigate the exercise-induced reduction in swimming capacity (Fig. 1*E-F*). Thus, our exercise regimen was within a physiological range that did not cause irreversible harm. Our findings also revealed that the mitochondrial network is remodeled during a cycle of exercise and recovery, with initial fragmentation followed by fusion and network reorganization that parallels the recovery in exercise performance.

We also investigated how aging affects these responses to exercise. Compared to day 1 adults, adults at days 5 and 10 displayed a decline in performance throughout the swimming period, including more rapid fatigue, but partially recovered to their respective 0h baselines (Fig. 1*E-F*). These older animals also underwent a cycle of mitochondrial fragmentation and fusion, although the extent to which this remodeling occurred during the cycle was reduced as the network fragmentation increased with age (Fig. 1*D*). Thus, aging animals also responded to exercise with parallel cycles of performance decline and recovery, and mitochondrial fragmentation and network reorganization, even as fitness capacity and network integrity each declined (Fig. 1*G*).

### Mitochondrial fission and fusion are required for fitness capacity and exercise benefits

Our findings suggested that mitochondrial fission and fusion might be important for fitness capacity. Mitochondrial fission depends upon DRP-1 (dynamin related protein 1), and fusion upon FZO-1 (ortholog of mitofusins MFN1 and MFN2, responsible for fusing the outer membrane) and EAT-3 (ortholog of optical atrophy 1, which fuses the inner membrane) (Fig. 2*A*) (40, 41). Throughout their lifespan, worms in which either fission (*drp-1*) or fusion (*fzo-1*) is impaired displayed a substantial and sustained decline in fitness capacity compared to WT (Fig. 2*B*). Overexpression of these regulators also impaired fitness capacity (*SI Appendix*, Fig. S1A), and loss of their function prevented fitness capacity recovery during a cycle of exercise followed by rest (day 1, Fig. 2*C* and *SI Appendix*, Fig. S1B-C). Together, this suggests that fitness capacity depends upon optimal mitochondrial dynamics.

**Figure 2.**
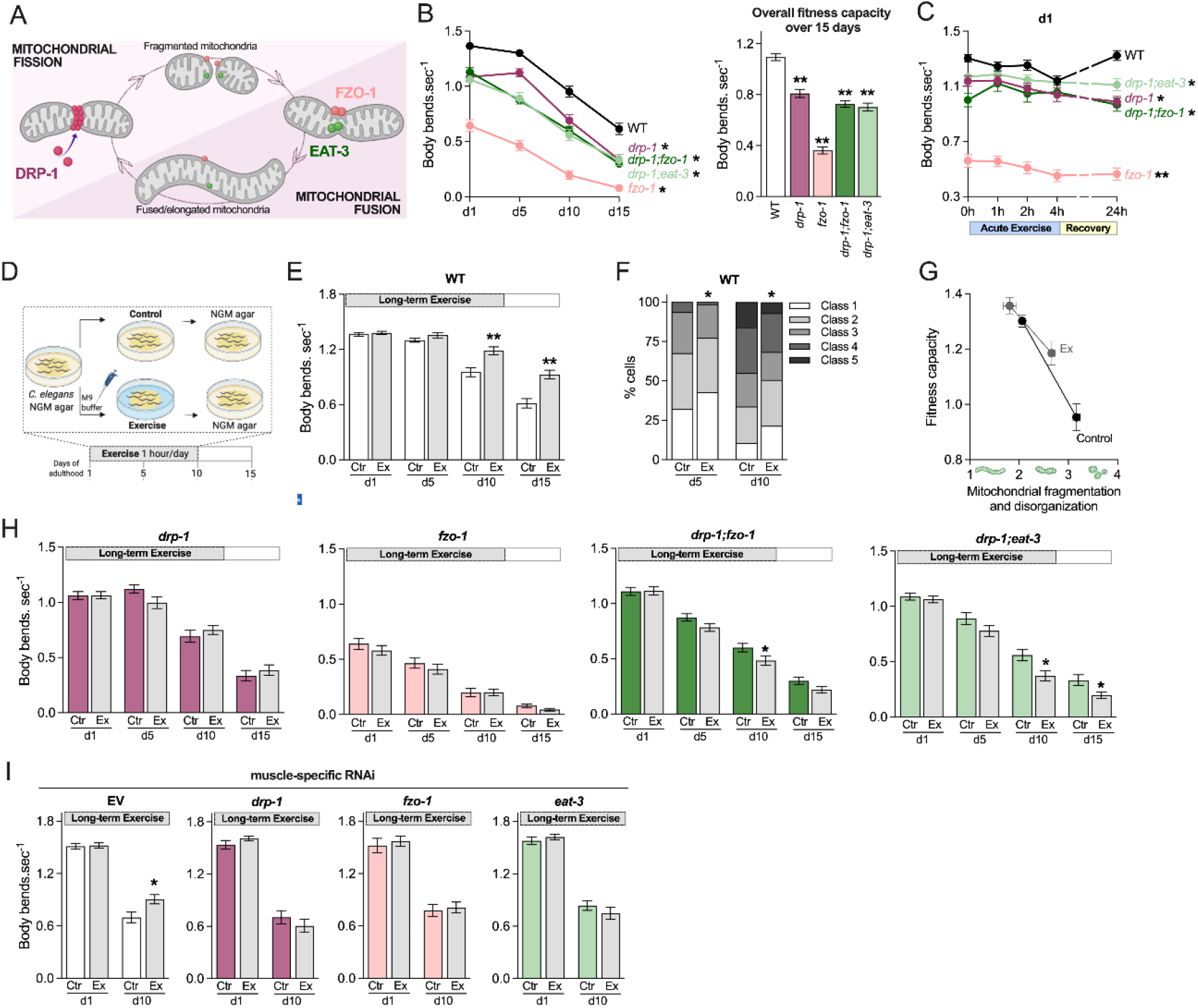
Mitochondrial fission and fusion are required for exercise-induced benefits during aging. (A) Simplified model for mitochondrial fusion and fission: mitochondrial fragmentation requires recruitment of cytosolic DRP-1 to the organelle, which oligomerizes and constricts the mitochondria into 2 daughters. The outer mitochondrial membrane fuses through interaction between FZO-1 of two opposing mitochondria, while EAT-3 drives inner mitochondrial membrane fusion. (B) Fitness capacity decay and overall fitness capacity (average of d1, d5, d10 and d15) of WT compared to mitochondrial dynamics mutants *drp-1(tm1108), fzo-1(tm1133), drp-1(tm1108);fzo-1(tm1133)* and *drp-1(tm1108);eat-3(ad426)* during aging. (C) Fitness capacity of WT and mitochondrial dynamics mutants *drp-1(tm1108), fzo-1(tm1133), drp-1(tm1108);fzo-1(tm1133)* and *drp-1(tm1108);eat-3(ad426)* submitted to acute exercise on day 1 of adulthood. (D) Long-term exercise: worms maintained at 20°C were submitted to 1 hour of exercise per day for 10 days, starting at the onset of adulthood (day 1). (E) Fitness capacity and (F) mitochondrial morphology in body wall muscle cells of WT worms submitted to long-term exercise. (G) Correlation between fitness capacity and mitochondrial morphology of control and long-term exercise-trained worms. (H) Fitness capacity of mitochondrial dynamics mutants *drp-1(tm1108), fzo-1(tm1133), drp-1(tm1108);fzo-1(tm1133)* and *drp-1(tm1108);eat-3(ad426)* submitted to long-term exercise. (I) Fitness capacity of muscle-specific RNAi strain *sid-1(qt-9);myo-3*p::sid-1 fed empty vector (EV) control or *drp-1, fzo-1* and *eat-3* RNAi from L4 stage, and submitted to long-term exercise. Data are presented as mean ± SEM. *p <0.05 and **p <0.001 vs. WT or Ctr. Detailed statistical analyses, numbers of biological replicates, and sample size are described in *SI Appendix*, Table S1.

In *C. elegans*, genetic inhibition of mitochondrial fission or fusion prevents some interventions from extending lifespan (12, 27). Remarkably, however, simultaneous disruption of fission and fusion can rescue these defects in some cases and even extends lifespan on its own, by preventing the mitochondrial fragmentation and some metabolic effects associated with aging (12). These last findings made it important to investigate how simultaneous prevention of fission and fusion affects fitness capacity. In striking contrast to its effects on lifespan, preventing mitochondrial fission and fusion (*drp-1;fzo-1* and *drp-1;eat-3*) failed to restore fitness capacity to WT levels, although ablation of fission ameliorated some effects of fusion loss (*drp-1;fzo-1* vs *fzo-1*) (Fig. 2*B-C*; *SI Appendix*, Fig. S1B-C). We conclude that maintenance of dynamic mitochondrial network remodeling *per se* is critical for fitness capacity.

Next, we investigated whether long-term exercise training might counteract the progressive loss of fitness capacity and mitochondrial connectiveness that occurs during aging, and whether mitochondrial dynamics is involved. We allowed WT animals to swim for one hour per day for 10 consecutive days, starting at the onset of adulthood (experimental day 1, Fig. 2*D*). This exercise regimen significantly improved fitness capacity at day 10, a benefit that was maintained for at least 5 days after exercise ceased (Fig. 2*E*), and mitigated the muscle mitochondrial fragmentation/disorganization seen during aging (Fig. 2*F*). Mitochondrial fragmentation and fitness capacity were also tightly and inversely correlated under these conditions (Fig. 2*G*). Disruption of mitochondrial fission (*drp-1*) or fusion (*fzo-1*) abrogated these long-term exercise benefits, and long-term exercise impaired fitness capacity when both fission and fusion were disrupted simultaneously (*drp-1;fzo-1* and *drp-1;eat-3*; Fig. 2*H*). Muscle-specific knockdown of these mitochondrial dynamics genes also abrogated the benefit of exercise (Fig. 2*I*). Taken together, our results indicate that exercise training delays the aging-associated decline in fitness capacity, dependent upon maintenance of mitochondrial fission and fusion in body-wall muscle.

### AMPK enhances fitness capacity through mitochondrial dynamics

Given that increases in mitochondrial fusion and density are associated with increased lifespan in *C. elegans* (12, 27, 42), we tested whether interventions that extend lifespan might in general improve exercise capacity during aging. We assessed fitness capacity in long-lived animals that are subject to mild mitochondrial dysfunction (*isp-1* and *nuo-6*) (43), reduced insulin/IGF-1 signaling (rIIS)(*daf-2*) (24), a dietary restriction (DR)-like state (*eat-2)*(44), or increased AMPK activity (45, 46). In the last case, we examined animals in which the AMPKα2 catalytic subunit carrying a constitutively activating mutation is overexpressed transgenically (CA-AAK-2 (46)). rIIS and DR extend lifespan across metazoans and have been shown to improve various healthspan parameters (24, 47), and AMPK is of particular interest here because it is a master regulator of energy homeostasis during exercise (48) and promotes remodelling of mitochondrial morphology and metabolism (12, 49).

Despite increasing lifespan, the *isp-1, nuo-6, daf-2* and *eat-2* mutations impaired fitness capacity and performance in an exercise-recovery cycle (Fig. 3*A*; *SI Appendix*, Fig. S2A-D). Compared to WT, *isp-1, nuo-6*, and *daf-2* mutants also displayed reduced fitness capacity with aging (day 10, Fig. 3*A*). A long-term exercise protocol did not slow the aging-related decline in fitness capacity in *isp-1, nuo-6, daf-2* or *eat-2* animals (*SI Appendix*, Fig. S2E-F), in contrast to its beneficial effect on WT worms (Fig. 2*E*). Together, the data suggest that mechanisms that extend lifespan are not necessarily sufficient to confer exercise-associated benefits.

**Figure 3.**
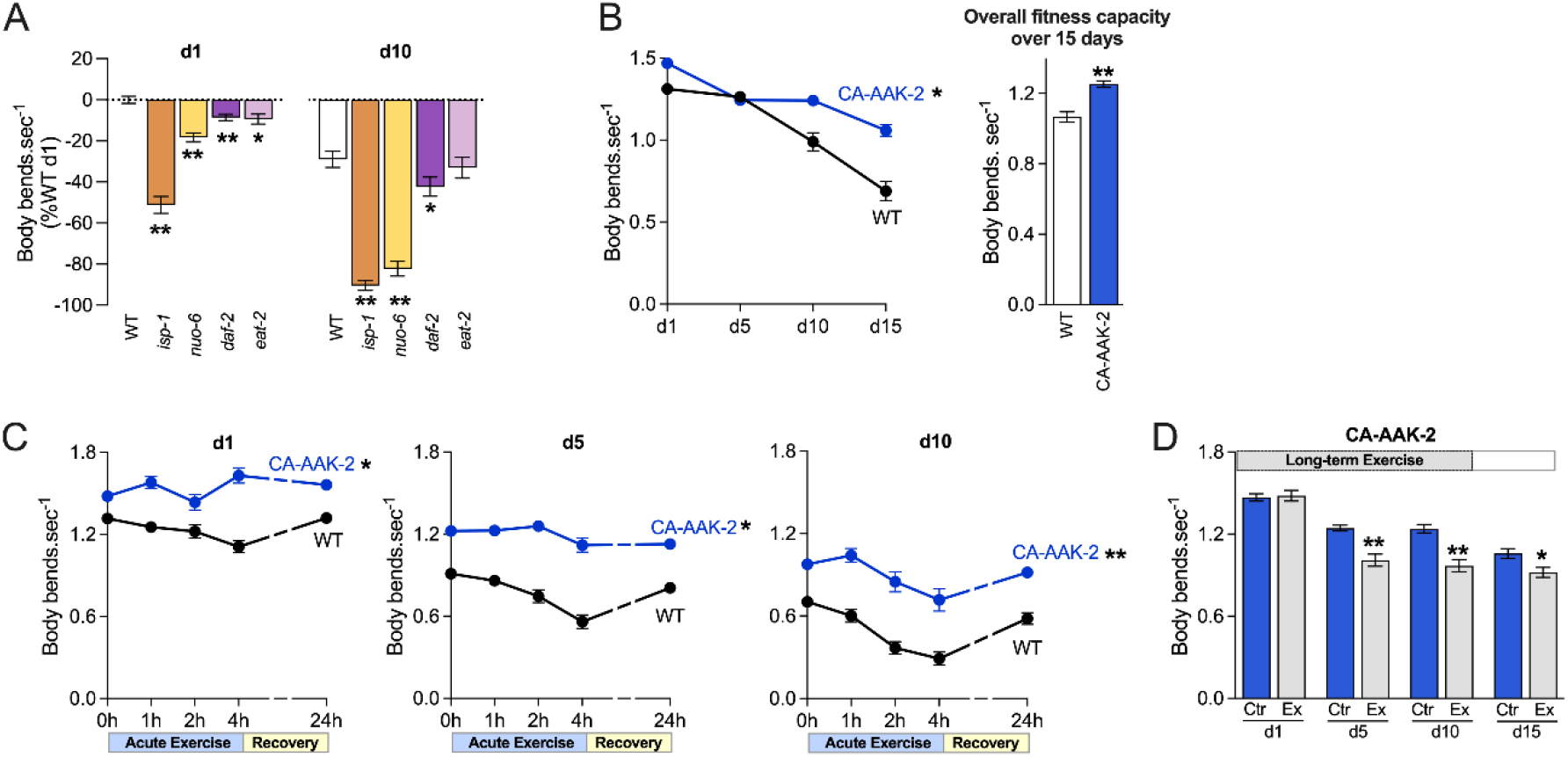
AMPK activation preserves fitness capacity with aging. (A) Fitness capacity decay of WT and the long-lived worms *isp-1(qm150), nuo-6(qm200), daf-2(e1370)* and *eat-2(ad1116)* with aging. (B) Fitness capacity decay and overall fitness capacity of WT and long-lived worms expressing constitutively active AMPK (CA-AAK-2) during aging. (C) Fitness capacity of WT and CA-AAK-2 worms submitted to acute exercise on days 1, 5 and 10 of adulthood. (D) Fitness capacity of CA-AAK-2 worms submitted to long-term exercise. Data are presented as mean ± SEM. *p <0.05 and **p <0.001 vs. WT or Ctr. Detailed statistical analyses, number of biological replicates, and sample size are described in *SI Appendix*, Table S1.

In striking contrast, animals expressing CA-AAK-2 exhibited improved fitness capacity and maintenance of exercise performance during aging (Fig. 3*B-C* and *SI Appendix*, Fig. S2G-H). Surprisingly, however, in CA-AAK-2 animals long-term exercise not only failed to further enhance fitness capacity during aging but was slightly detrimental (Fig. 3*D*). Thus, constitutive activation of AMPK dramatically improved fitness capacity with age but did not allow further benefit from exercise.

We investigated whether AMPK might be required for fitness capacity and exercise benefits. Compared to WT, AMPK-deficient mutants (*aak-2*) exhibited a sustained reduction in fitness capacity during aging that was lost at day 15 (Fig. 4*A*), as well as impairment of the exercise-recovery cycle (at day 1, Fig. 4*B* and *SI Appendix*, Fig. S3A-B). Finally, AMPK-deficient mutants did not benefit from long-term exercise over the course of their lifespan, and even exhibited reduced exercise capacity compared with non-exercised animals (Fig. 4*C*). Thus, not only does AMPK enhance fitness capacity during aging when constitutively activated (Fig. 3*B*), it is also required for normal exercise responsiveness and benefits throughout life.

**Figure 4.**
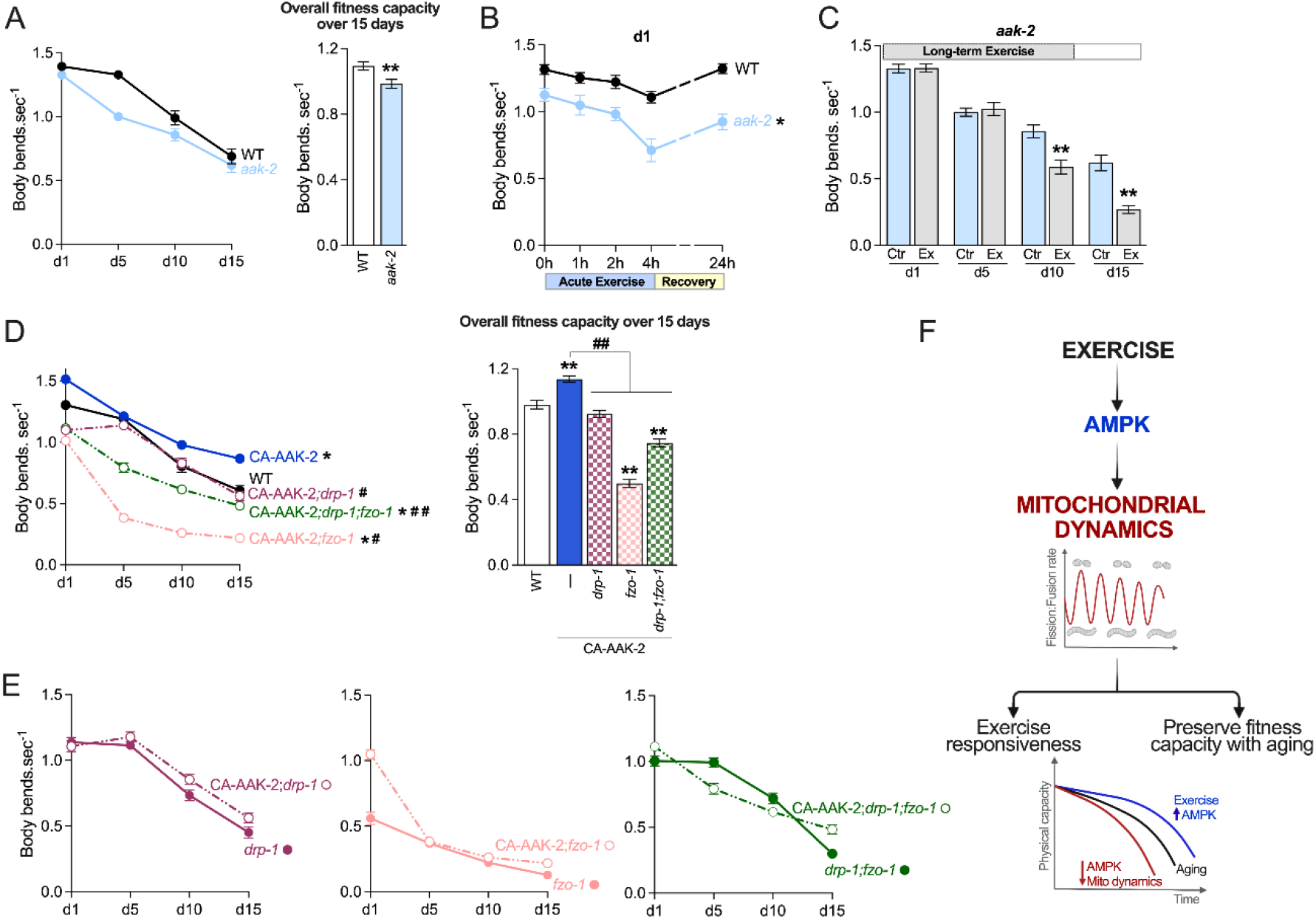
The AMPK-induced improvement in fitness capacity requires mitochondrial dynamics. (A) Fitness capacity decay and overall fitness capacity of WT and AMPK-deficient worms (*aak-2(gt33)*) during aging. (B) Fitness capacity of WT and *aak-2(gt33)* day-one adults subjected to an exercise-recovery cycle. (C) Fitness capacity of *aak-2(gt33)* worms subjected to long-term exercise. (D) Fitness capacity decay and overall fitness capacity of animals carrying the indicated mitochondrial dynamics mutations (*drp-1(tm1108), fzo-1(tm1133)* and *drp-1(tm1108);fzo-1(tm1133)*) in either the WT or CA-AAK-2 background during aging. CA-AAK-2 in the WT background is presented in blue. (E) Fitness capacity decay of the indicated strains during aging. Data are presented as mean ± SEM. *p <0.05 and **p <0.001 vs. WT or Ctr. #p <0.05 and ##p <0.001 vs. CA-AAK-2. (F) Working model for maintenance of exercise responsiveness and fitness capacity during aging: the beneficial effects of exercise are mediated through AMPK and mitochondrial dynamics. Detailed statistical analyses, number of biological replicates, and sample size are described in *SI Appendix*, Table S1.

Our findings raise the question of whether the salutary effects of increased AMPK activity on exercise performance during aging might depend upon mitochondrial dynamics. Supporting this idea, impairment of mitochondrial fission (*drp-1*), fusion (*fzo-1*), or both processes (*drp-1;fzo-1*) abrogated the CA-AAK2-induced improvements in fitness capacity during aging (Fig. 4*D*). In each case, the negative effects of fission or fusion loss on swimming performance were similar in CA-AAK-2 and WT animals (Fig. 4*E* and *SI Appendix*, Fig. S3C). Together, these findings indicate that mitochondrial dynamics is an essential effector of AMPK activity in promoting fitness capacity.

## Discussion

Despite extensive evidence that impairment of mitochondrial fission or fusion contributes to age-related diseases (50, 51), the role of mitochondrial dynamics in anti-aging interventions such as exercise has remained uncertain. Here, by taking advantage of tools available for *C. elegans* we demonstrated that mitochondrial fission and fusion are each required for both fitness capacity and its improvement by exercise training, and that mitochondrial network plasticity – the capacity for re-shaping the mitochondrial network between fused and fragmented states, is also crucial (Fig. 4*F*). Moreover, the only anti-aging intervention we identified that enhances fitness capacity during aging, AMPK activation, depends upon mitochondrial dynamics and plasticity for this benefit. Given that the mechanisms that mediate mitochondrial dynamics are evolutionarily conserved, it seems likely that the generally critical role for mitochondrial dynamics and plasticity we uncovered may be broadly applicable.

One important finding was that a single exercise session induces a cycle of fatigue and fitness capacity recovery that is paralleled by a cycle of mitochondrial fission and network rebuilding in body wall muscle. The extent to which these parameters change during the exercise/recovery cycle was dampened with age, in parallel to a decline in fitness capacity. By contrast, a regular exercise training regimen enhanced fitness capacity, delaying its inevitable decline during aging. Mitochondrial dynamics was critical for fitness capacity under each of these conditions, and its disruption by impairment of fission or fusion dramatically impaired fitness capacity and ablated the benefits of exercise.

Critically, “freezing” the mitochondrial network by simultaneous disruption of both fission and fusion similarly impaired fitness capacity, demonstrating that network plasticity is essential during the exercise/recovery cycle. This contrasts sharply to one effect of mitochondrial dynamics on aging, because simultaneous ablation of mitochondrial fission and fusion can extend lifespan, apparently by delaying the aging-induced degradation of the mitochondrial network and accompanying metabolic perturbations (12). Thus, while maintaining the mitochondrial network is sufficient to slow aging, maintaining the capacity for network remodeling in muscle seems to be critical for meeting the metabolic demands of exercise. This need for plasticity may explain why most anti-aging interventions we tested impaired fitness capacity during aging, because two of those interventions (rIIS and DR) maintain network structure during aging (12, 27). It is also consistent with our findings that simultaneous ablation of fission and fusion blocks lifespan extension from intermittent fasting, which like exercise induces cycles of mitochondrial network fission and recovery (12).

It is striking that constitutive AMPK activation uniquely enhanced fitness capacity during aging, benefitting the animal comparably to an exercise regimen. In mammals AMPK acts as an key signaling molecule in regulating mitochondrial homeostasis during exercise (52-54), although its role in mediating exercise benefits has remained inconclusive (55). Here we demonstrated in *C. elegans* that AMPK is required for both baseline fitness capacity and exercise-induced benefits, and that AMPK activation enhances fitness capacity dependent upon mitochondrial dynamics. Together, our results suggest that exercise benefits may be conferred through a linear pathway involving AMPK regulation of mitochondrial dynamics (Fig. 4*F*). Interestingly, exercise training did not improve fitness capacity in the setting of constitutive AMPK activation. This might indicate a maximum limit in fitness capacity, but it is also consistent with the idea that AMPK activation might have acted as a surrogate for exercise training in conferring these benefits. The anti-diabetic drug metformin has many effects on metabolism, including stimulating AMPK indirectly (56). It may be consistent with our results that metformin not only extends *C. elegans* lifespan but also delays the decline in swimming capacity with aging (57).

An important goal of the aging field is to identify interventions that not only extend lifespan, but also enhance important parameters of health (1). Our approach suggests that a multipronged approach may be necessary, given the unique demands of fitness capacity and its maintenance through exercise. In aging humans, a decline in muscle function and exercise tolerance is a major concern that leads to substantial morbidity (58). Our data point towards AMPK and mitochondrial dynamics as potentially fruitful intervention points for forestalling this decline, most likely along with other aspects of aging. Considering the clinical implications of coordinating lifespan with healthspan, it will be of great interest to determine how mitochondrial network plasticity influences fitness capacity, along with longevity and aging-associated disease, in humans.

## Materials and Methods

### Strains and maintenance of *C. elegans*

*C. elegans* strains used in this study are listed *in SI Appendix*, Table S2. Nematodes were grown and maintained at 20°C on standard nematode growth media (NGM) agar plates seeded with live *Escherichia coli* (*E. coli*). *E. coli* (OP50-1) was cultured overnight in LB medium containing 10 mg/L streptomycin at 37°C. RNAi experiments were performed by using tetracycline–resistant *E. coli* (HT115) carrying dsRNA against the genes *drp-1, fzo-1* and *eat-3*, or an empty vector control (pL4440). RNAi cultures were grown overnight at 37°C in LB medium containing 50 mg/mL carbenicillin and dsRNA expression was induced by the addition of 0.2 g/L IPTG prior to seed onto NGM agar plates containing 50 mg/mL carbenicillin and 0.2 g/L IPTG. Synchronized populations of L1 animals were obtained by hypochlorite treatment (59), then allowed to develop at 20°C on seeded NGM agar plates. Whenever experiments were conducted over lifespan, 20 mg/L FUdR was added at the L4 stage to prevent hatching.

### Fitness capacity

Fitness capacity was measured by calculating the body bends per second of worms in liquid environment as previously described (17) with minor modifications. Briefly, we transferred worms to 96-well plates containing M9 buffer (1 worm per well) and immediately recorded a 30-seconds video at a rate of 15 frames per second using a stereomicroscope (Optika SMZ-4) coupled with a device camera. Recorded images were analyzed using the ImageJ plugin Worm-tracker (wrMTrck) (60). At least 15 animals were recorded per biological replicate.

### Mitochondrial morphology

Mitochondrial network morphology was detected using strains expressing green fluorescent protein (GFP) targeted to the outer mitochondrial membrane specifically in the body wall muscle (zcIs14[*myo-3*::GFP(mito)]). Worms were anesthetized in Tetramisole 0.2 mg.ml^−1^, mounted on 2% agarose pads on glass slides, and subsequently imaged on Zeiss Axio Imager M2 fluorescence microscope with Axiocam HRC camera. Muscle mitochondria were analyzed in cells midway between the pharynx and vulva, or vulva and tail. Qualitative assessment of mitochondrial morphology was made by scoring cells based on five classes as previously described by Laranjeiro et. al (29). These categories reflect a progressive increase in fragmentation and disorganization from class 1 (tubular and interconnected mitochondrial network) to class 5 (reduced number of fragmented mitochondria) (Fig. 1*B*). Images were taken of > 35 muscle cells from at least 15 worms per biological replicate. All analyses were conducted by a single observer, blinded to animal’s identity.

### Exercise protocols

#### Acute exercise

Acute exercise was mimicked by putting worms to swim for 4 hours. Briefly, seeded NGM agar plates were flooded with M9 buffer. After 4 hours, worms were washed off with M9 buffer into 15 mL conical tubes, centrifuged at 700 rpm for 1 min, transferred to seeded NGM agar plates and allowed to rest for 24 hours at 20°C (recovery period) (Fig. 1*C*). Because some of the swimming benefits have been recently attributed to food transient deprivation (30), the exercise session was performed in the presence of food by adding M9 buffer to NGM agar plates containing bacteria.

#### Long-term exercise

Long-term exercise was performed by allowing worms to swim 1 hour per day for 10 days, starting at the onset of adulthood (day 1) (Fig. 2*D*). Briefly, worm strains were divided into control and exercise groups and maintained at 20°C on separated 100 mm seeded NGM agar plates. Exercise group plates were flooded with 10 mL of M9 buffer. After 1 hour, worms were washed off with M9 buffer into 15 mL conical tubes, centrifuged at 700 rpm for 1 min and transferred to seeded NGM agar plates using a glass Pasteur pipette to minimize nematode loss. Control animals were also transferred to seeded NGM agar plates using the same method. This procedure was repeated daily for the next 10 days.

### Statistics

Data obtained in this study are presented as mean ± standard error of the mean (SEM) and are compiled from multiple trials. Chi-square test was used to compare the distribution of muscle mitochondrial morphology into multiple categories (Fig. 1*B*, Fig. 1*D*, and Fig. 2*F*). Linear regression was used to assess the association between variables in Fig. 1*G* and Fig. 2*G*. For all other assays, two-tailed Student’s t test was used for comparison between 2 groups. GraphPad Prism was used for statistical analyses and statistical significance was considered achieved when the value of *p* was < 0.05. Detailed statistical analyses, biological replicates (independent population of worms tested on a different day) and sample size are described in *SI Appendix*, Table S1.

## Data availability

All study data are included in the article and/or *SI Appendix*.

## Acknowledgments

This work was supported by Fundação de Amparo à Pesquisa do Estado de São Paulo (FAPESP) 2013/07937-8, 2015/22814-5, 2017/16694-2 and 2019/25049-9; Conselho Nacional de Pesquisa e Desenvolvimento – Brasil (CNPq) 303281/2015-4 and 407306/2013-7; Coordenação de Aperfeiçoamento de Pessoal de Nível Superior – Brasil (CAPES) Finance Code 001 and Instituto Nacional de Ciência e Tecnologia and Centro de Pesquisa e Desenvolvimento de Processos Redox em Biomedicina to J.C.B.F.; National Institutes of Health (NIH) R35 GM122610, R01 AG054215 to T.K.B., the Joslin Diabetes Center P30 DK036836, and R01 GM121756 to J.V.R.; FAPESP postdoctoral fellowships 2017/16540-5 and 2019/18444-9 to J.C.C., and 2016/09611-0 and 2019/07221-9 to L.H.M.B. We thank William B. Mair (Harvard T.H. Chan School of Public Health) and Malena Hansen (Sanford Burnham Prebys Medical Discovery Institute) for providing some of the worm strains used in this study. Other strains were provided by the CGC, which is funded by the NIH (P40 OD010440).

## Author Contributions

J.C.C., L.H.M.B., T.K.B. and J.C.B.F. designed research; J.C.C. and L.H.M.B. performed research and analyzed data; A.T., A.V.B. and J.V.R. contributed new reagents/analytic tools; J.V.R., T.K.B. and J.C.B.F. supervision; J.C.C., T.K.B. and J.C.B.F. wrote the manuscript, and J.C.C., L.H.M.B., A.T., A.V.B., J.V.R., T.K.B. and J.C.B.F. reviewed and edited the manuscript.

## References

1. J. Campisi et al., From discoveries in ageing research to therapeutics for healthy ageing. Nature 571, 183–192 (2019).

2. C. Lopez-Otin, M. A. Blasco, L. Partridge, M. Serrano, G. Kroemer, The hallmarks of aging. Cell 153, 1194–1217 (2013).

3. S. G. Regmi, S. G. Rolland, B. Conradt, Age-dependent changes in mitochondrial morphology and volume are not predictors of lifespan. Aging (Albany NY) 6, 118–130 (2014).

4. J. C. B. Ferreira et al., A selective inhibitor of mitofusin 1-betaIIPKC association improves heart failure outcome in rats. Nat Commun 10, 329 (2019).

5. N. Taguchi, N. Ishihara, A. Jofuku, T. Oka, K. Mihara, Mitotic phosphorylation of dynamin-related GTPase Drp1 participates in mitochondrial fission. J Biol Chem 282, 11521–11529 (2007).

6. G. Twig et al., Fission and selective fusion govern mitochondrial segregation and elimination by autophagy. EMBO J 27, 433–446 (2008).

7. H. Chen, A. Chomyn, D. C. Chan, Disruption of fusion results in mitochondrial heterogeneity and dysfunction. J Biol Chem 280, 26185–26192 (2005).

8. O. S. Shirihai, M. Song, G. W. Dorn, 2nd, How mitochondrial dynamism orchestrates mitophagy. Circ Res 116, 1835–1849 (2015).

9. D. Sebastian et al., Mfn2 deficiency links age-related sarcopenia and impaired autophagy to activation of an adaptive mitophagy pathway. EMBO J 35, 1677–1693 (2016).

10. J. J. Byrne et al., Disruption of mitochondrial dynamics affects behaviour and lifespan in Caenorhabditis elegans. Cell Mol Life Sci 76, 1967–1985 (2019).

11. C. Tezze et al., Age-Associated Loss of OPA1 in Muscle Impacts Muscle Mass, Metabolic Homeostasis, Systemic Inflammation, and Epithelial Senescence. Cell Metab 25, 1374–1389 e1376 (2017).

12. H. J. Weir et al., Dietary Restriction and AMPK Increase Lifespan via Mitochondrial Network and Peroxisome Remodeling. Cell Metab 26, 884–896 e885 (2017).

13. Y. Wang, J. Nie, G. Ferrari, J. P. Rey-Lopez, L. F. M. Rezende, Association of Physical Activity Intensity With Mortality: A National Cohort Study of 403681 US Adults. JAMA Intern Med 181, 203–211 (2021).

14. V. Gremeaux et al., Exercise and longevity. Maturitas 73, 312–317 (2012).

15. J. R. Beard et al., The World report on ageing and health: a policy framework for healthy ageing. Lancet 387, 2145–2154 (2016).

16. S. Studenski et al., Gait speed and survival in older adults. JAMA 305, 50–58 (2011).

17. J. H. Hahm et al., C. elegans maximum velocity correlates with healthspan and is maintained in worms with an insulin receptor mutation. Nat Commun 6, 8919 (2015).

18. U. Wisloff et al., Cardiovascular risk factors emerge after artificial selection for low aerobic capacity. Science 307, 418–420 (2005).

19. J. A. Hawley, M. Hargreaves, M. J. Joyner, J. R. Zierath, Integrative biology of exercise. Cell 159, 738–749 (2014).

20. J. C. Campos et al., Exercise reestablishes autophagic flux and mitochondrial quality control in heart failure. Autophagy 13, 1304–1317 (2017).

21. M. Coronado et al., Physiological Mitochondrial Fragmentation Is a Normal Cardiac Adaptation to Increased Energy Demand. Circ Res 122, 282–295 (2018).

22. T. M. Moore et al., The impact of exercise on mitochondrial dynamics and the role of Drp1 in exercise performance and training adaptations in skeletal muscle. Mol Metab 21, 51–67 (2019).

23. M. B. Bell, Z. Bush, G. R. McGinnis, G. C. Rowe, Adult skeletal muscle deletion of Mitofusin 1 and 2 impedes exercise performance and training capacity. J Appl Physiol (1985) 126, 341–353 (2019).

24. C. J. Kenyon, The genetics of ageing. Nature 464, 504–512 (2010).

25. H. S. Chuang, W. J. Kuo, C. L. Lee, I. H. Chu, C. S. Chen, Exercise in an electrotactic flow chamber ameliorates age-related degeneration in Caenorhabditis elegans. Sci Rep 6, 28064 (2016).

26. E. Teo et al., A novel vibration-induced exercise paradigm improves fitness and lipid metabolism of Caenorhabditis elegans. Sci Rep 8, 9420 (2018).

27. S. N. Chaudhari, E. T. Kipreos, Increased mitochondrial fusion allows the survival of older animals in diverse C. elegans longevity pathways. Nat Commun 8, 182 (2017).

28. R. Laranjeiro, G. Harinath, D. Burke, B. P. Braeckman, M. Driscoll, Single swim sessions in C. elegans induce key features of mammalian exercise. BMC Biol 15, 30 (2017).

29. R. Laranjeiro et al., Swim exercise in Caenorhabditis elegans extends neuromuscular and gut healthspan, enhances learning ability, and protects against neurodegeneration. Proc Natl Acad Sci U S A 116, 23829–23839 (2019).

30. J. H. Hartman et al., Swimming Exercise and Transient Food Deprivation in Caenorhabditis elegans Promote Mitochondrial Maintenance and Protect Against Chemical-Induced Mitotoxicity. Sci Rep 8, 8359 (2018).

31. A. Sujkowski, B. Bazzell, K. Carpenter, R. Arking, R. J. Wessells, Endurance exercise and selective breeding for longevity extend Drosophila healthspan by overlapping mechanisms. Aging (Albany NY) 7, 535–552 (2015).

32. N. Piazza, B. Gosangi, S. Devilla, R. Arking, R. Wessells, Exercise-training in young Drosophila melanogaster reduces age-related decline in mobility and cardiac performance. PLoS One 4, e5886 (2009).

33. S. Studenski et al., Physical performance measures in the clinical setting. J Am Geriatr Soc 51, 314–322 (2003).

34. H. C. Jiang et al., Neural activity and CaMKII protect mitochondria from fragmentation in aging Caenorhabditis elegans neurons. Proc Natl Acad Sci U S A 112, 8768–8773 (2015).

35. J. P. Leduc-Gaudet et al., Mitochondrial morphology is altered in atrophied skeletal muscle of aged mice. Oncotarget 6, 17923–17937 (2015).

36. A. Rana et al., Promoting Drp1-mediated mitochondrial fission in midlife prolongs healthy lifespan of Drosophila melanogaster. Nat Commun 8, 448 (2017).

37. M. Hargreaves, L. L. Spriet, Skeletal muscle energy metabolism during exercise. Nat Metab 2, 817–828 (2020).

38. M. Liesa, O. S. Shirihai, Mitochondrial dynamics in the regulation of nutrient utilization and energy expenditure. Cell Metab 17, 491–506 (2013).

39. P. Mishra, D. C. Chan, Metabolic regulation of mitochondrial dynamics. J Cell Biol 212, 379–387 (2016).

40. A. M. Labrousse, M. D. Zappaterra, D. A. Rube, A. M. van der Bliek, C. elegans dynamin-related protein DRP-1 controls severing of the mitochondrial outer membrane. Mol Cell 4, 815–826 (1999).

41. B. P. Head, M. Zulaika, S. Ryazantsev, A. M. van der Bliek, A novel mitochondrial outer membrane protein, MOMA-1, that affects cristae morphology in Caenorhabditis elegans. Mol Biol Cell 22, 831–841 (2011).

42. N. S. Morsci, D. H. Hall, M. Driscoll, Z. H. Sheng, Age-Related Phasic Patterns of Mitochondrial Maintenance in Adult Caenorhabditis elegans Neurons. J Neurosci 36, 1373–1385 (2016).

43. W. Yang, S. Hekimi, Two modes of mitochondrial dysfunction lead independently to lifespan extension in Caenorhabditis elegans. Aging Cell 9, 433–447 (2010).

44. B. Lakowski, S. Hekimi, The genetics of caloric restriction in Caenorhabditis elegans. Proc Natl Acad Sci U S A 95, 13091–13096 (1998).

45. J. Apfeld, G. O’Connor, T. McDonagh, P. S. DiStefano, R. Curtis, The AMP-activated protein kinase AAK-2 links energy levels and insulin-like signals to lifespan in C. elegans. Genes Dev 18, 3004–3009 (2004).

46. W. Mair et al., Lifespan extension induced by AMPK and calcineurin is mediated by CRTC-1 and CREB. Nature 470, 404–408 (2011).

47. L. Fontana, L. Partridge, Promoting health and longevity through diet: from model organisms to humans. Cell 161, 106–118 (2015).

48. S. Herzig, R. J. Shaw, AMPK: guardian of metabolism and mitochondrial homeostasis. Nat Rev Mol Cell Biol 19, 121–135 (2018).

49. K. Burkewitz et al., Neuronal CRTC-1 governs systemic mitochondrial metabolism and lifespan via a catecholamine signal. Cell 160, 842–855 (2015).

50. B. N. Whitley, E. A. Engelhart, S. Hoppins, Mitochondrial dynamics and their potential as a therapeutic target. Mitochondrion 49, 269–283 (2019).

51. L. H. M. Bozi, J. C. Campos, V. O. Zambelli, N. D. Ferreira, J. C. B. Ferreira, Mitochondrially-targeted treatment strategies. Mol Aspects Med 71, 100836 (2020).

52. N. J. Hoffman et al., Global Phosphoproteomic Analysis of Human Skeletal Muscle Reveals a Network of Exercise-Regulated Kinases and AMPK Substrates. Cell Metab 22, 922–935 (2015).

53. S. Jager, C. Handschin, J. St-Pierre, B. M. Spiegelman, AMP-activated protein kinase (AMPK) action in skeletal muscle via direct phosphorylation of PGC-1alpha. Proc Natl Acad Sci U S A 104, 12017–12022 (2007).

54. R. C. Laker et al., Ampk phosphorylation of Ulk1 is required for targeting of mitochondria to lysosomes in exercise-induced mitophagy. Nat Commun 8, 548 (2017).

55. B. Viollet et al., AMPK: Lessons from transgenic and knockout animals. Front Biosci (Landmark Ed) 14, 19–44 (2009).

56. J. I. Castillo-Quan, T. K. Blackwell, Metformin: Restraining Nucleocytoplasmic Shuttling to Fight Cancer and Aging. Cell 167, 1670–1671 (2016).

57. B. Onken et al., Metformin treatment of diverse Caenorhabditis species reveals the importance of genetic background in longevity and healthspan extension outcomes. Aging Cell 21, e13488 (2022).

58. D. R. Seals, J. N. Justice, T. J. LaRocca, Physiological geroscience: targeting function to increase healthspan and achieve optimal longevity. J Physiol 594, 2001–2024 (2016).

59. M. Porta-de-la-Riva, L. Fontrodona, A. Villanueva, J. Ceron, Basic Caenorhabditis elegans methods: synchronization and observation. J Vis Exp 10.3791/4019, e4019 (2012).

60. C. I. Nussbaum-Krammer, M. F. Neto, R. M. Brielmann, J. S. Pedersen, R. I. Morimoto, Investigating the spreading and toxicity of prion-like proteins using the metazoan model organism C. elegans. J Vis Exp 10.3791/52321, 52321 (2015).

