## Supplemental Information for "Exercise preserves fitness capacity during aging through AMPK and mitochondrial dynamics"

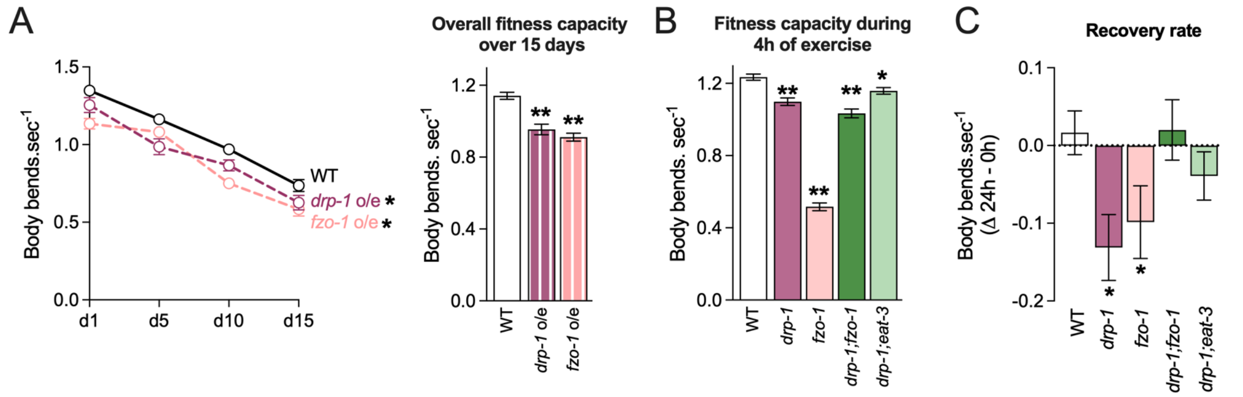


**Fig. S1. Impaired mitochondrial fission or fusion compromises fitness capacity (related to Fig. 2).** (A) Fitness capacity decay and overall fitness capacity (average of d1, d5, d10 and d15) of worms overexpressing mitochondrial dynamics genes *drp-1* o/e and *fzo-1* o/e with aging. (B) Fitness capacity during 4h of exercise and (C) recovery rate of WT and mitochondrial dynamics mutants *drp-1(tm1108), fzo-1(tm1133), drp-1(tm1108);fzo-1(tm1133)* and *drp-1(tm1108);eat-3(ad426)* submitted to acute exercise on day 1 of adulthood. Data are presented as mean ± SEM. *p <0.05 and **p <0.001 vs. WT. Detailed statistical analyses, number of biological replicates and sample size are described in *SI Appendix*, Table S1.

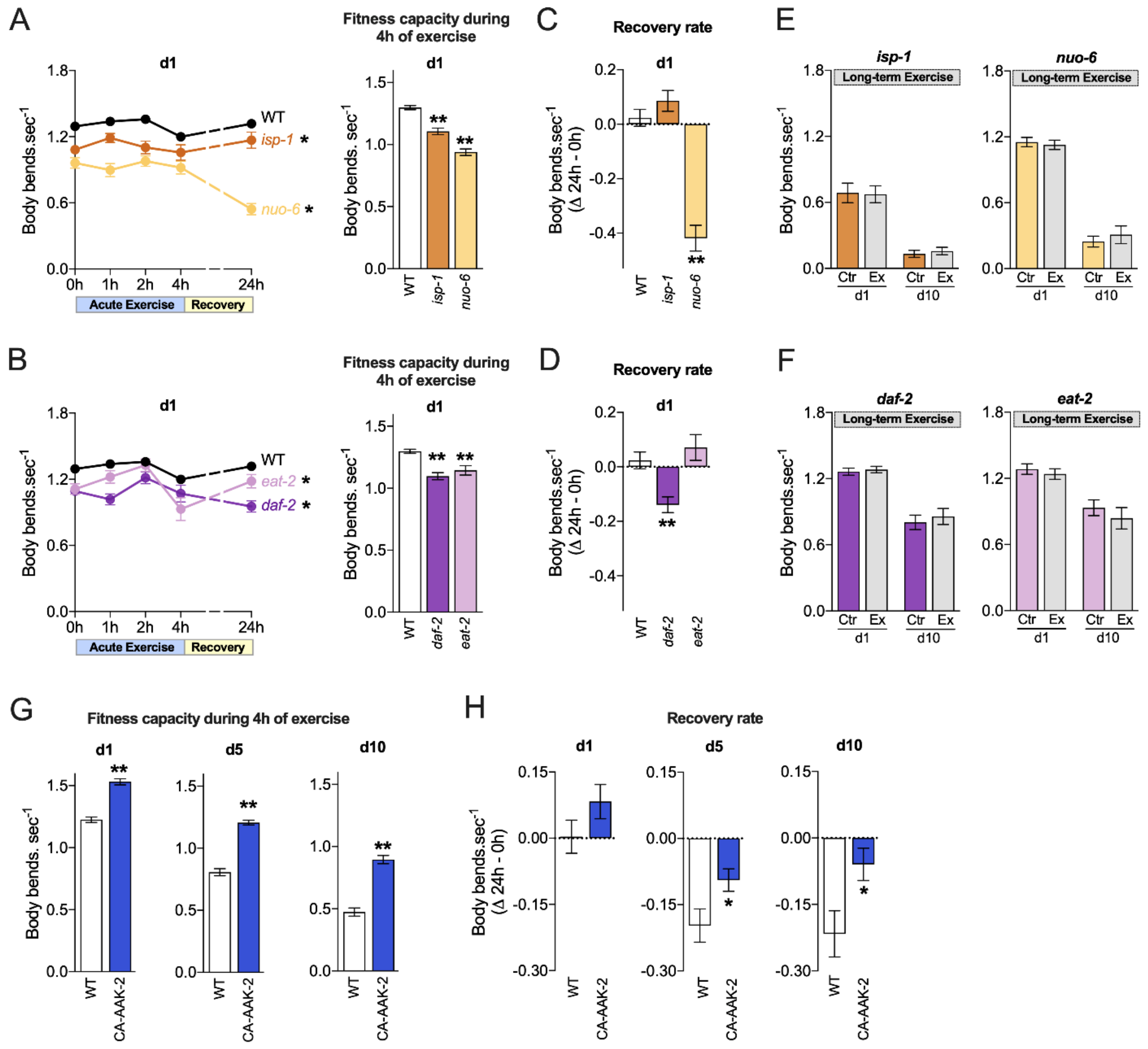


**Fig. S2. Long-lived mutants isp-1, nuo-6, daf-2 and eat-2 have decreased fitness capacity and exhibit no beneficial effects of exercise (related to Fig. 3).** (A, B) Fitness capacity during 4h of exercise and (C, D) recovery rate of WT, *isp-1(qm150), nuo-6(qm200), daf-2(e1370)* and *eat-2(ad1116)* worms submitted to acute exercise on day 1 of adulthood. (E, F) Fitness capacity of *isp-1(qm150), nuo-6(qm200), daf-2(e1370)* and *eat-2(ad1116)* worms submitted to long-term exercise. (G) Fitness capacity over 4h of exercise and (H) recovery rate of WT and CA-AAK-2 worms submitted to acute exercise on days 1, 5 and 10 of adulthood. Data are presented as mean ± SEM. *p <0.05 and **p <0.001 vs. WT. Detailed statistical analyses, number of biological replicates and sample size are described in *SI Appendix*, Table S1.

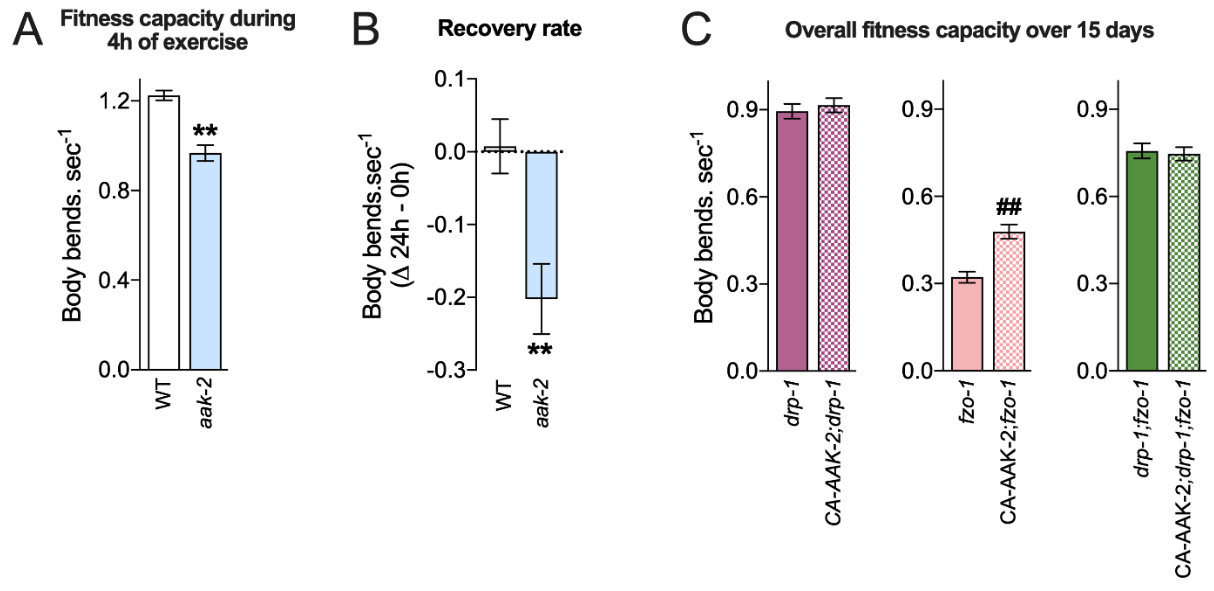


**Fig. S3. AMPK is required for exercise-induced benefits (related to Fig. 4).** (A) Fitness capacity during 4h of exercise and (B) recovery rate of WT and *aak-2(gt33)* worms submitted to acute exercise on day 1 of adulthood. (C) Overall fitness capacity (average of d1, d5, d10 and d15) of mitochondrial dynamics mutants *drp-1(tm1108), fzo-1(tm1133)* and *drp-1(tm1108);fzo-1(tm1133)* in the absence or presence of CA-AAK-2 with aging. Data are presented as mean ± SEM. **p <0.001 vs. WT and ##p <0.001 vs. *fzo-1(tm1133)*. Detailed statistical analyses, number of biological replicates and sample size are described in *SI Appendix*, Table S1.

**Table S1. Biological replicates, sample size and statistical analyses.**

| Figure (Replicates) | Strain - Group | Sample size | Statistical analyses | *p* value |
|---|---|---|---|---|
| 1A (3) | zcIs14[*myo-3*::GFP(mito)] - d1 | 56 | | |
| | zcIs14[*myo-3*::GFP(mito)] - d5 | 67 | two-tailed Student's t vs. d1 | <.0001 |
| | zcIs14[*myo-3*::GFP(mito)] - d10 | 66 | two-tailed Student's t vs. d1 | <.0001 |
| | zcIs14[*myo-3*::GFP(mito)] - d15 | 45 | two-tailed Student's t vs. d1 | <.0001 |
| 1B (3) | zcIs14[*myo-3*::GFP(mito)] - d1 | 126 | | |
| | zcIs14[*myo-3*::GFP(mito)] - d5 | 173 | chi-square vs. d1 | <.0001 |
| | zcIs14[*myo-3*::GFP(mito)] - d10 | 151 | chi-square vs. d1 | <.0001 |
| 1D (3) | zcIs14[*myo-3*::GFP(mito)] - d1 0h | 126 | | |
| | zcIs14[*myo-3*::GFP(mito)] - d1 1h Exercise | 138 | chi-square vs. d1 0h | <.0001 |
| | zcIs14[*myo-3*::GFP(mito)] - d1 2h Exercise | 149 | chi-square vs. d1 0h | <.0001 |
| | zcIs14[*myo-3*::GFP(mito)] - d1 4h Exercise | 173 | chi-square vs. d1 0h | <.0001 |
| | zcIs14[*myo-3*::GFP(mito)] - d1 24h Recovery | 150 | chi-square vs. d1 0h | 0.0001 |
| | | | chi-square vs. d1 4h | <.0001 |
| 1D (3) | zcIs14[*myo-3*::GFP(mito)] - d5 0h | 156 | | |
| | zcIs14[*myo-3*::GFP(mito)] - d5 1h Exercise | 159 | chi-square vs. d5 0h | 0.3565 |

| | | | | |
|---|---|---|---|---|
| | zcIs14[*myo-3*::GFP(mito)] - d5 2h Exercise | 165 | chi-square vs. d5 0h | 0.0009 |
| | zcIs14[*myo-3*::GFP(mito)] - d5 4h Exercise | 154 | chi-square vs. d5 0h | <.0001 |
| | zcIs14[*myo-3*::GFP(mito)] - d5 24h Recovery | 156 | chi-square vs. d5 0h | 0.6592 |
| | | | chi-square vs. d5 4h | <.0001 |
| 1D (2) | zcIs14[*myo-3*::GFP(mito)] - d10 0h | 135 | | |
| | zcIs14[*myo-3*::GFP(mito)] - d10 1h Exercise | 74 | chi-square vs. d10 0h | 0.8980 |
| | zcIs14[*myo-3*::GFP(mito)] - d10 2h Exercise | 83 | chi-square vs. d10 0h | 0.9652 |
| | zcIs14[*myo-3*::GFP(mito)] - d10 4h Exercise | 86 | chi-square vs. d10 0h | 0.2928 |
| | zcIs14[*myo-3*::GFP(mito)] - d10 24h Recovery | 75 | chi-square vs. d10 0h | 0.1979 |
| | | | chi-square vs. d10 4h | 0.0135 |
| 1E (3) | zcIs14[*myo-3*::GFP(mito)] - d1 0h | 56 | | |
| | zcIs14*[myo-3*::GFP(mito)] - d1 1h Exercise | 55 | two-tailed Student's t vs. d1 0h | 0.1744 |
| | zcIs14[*myo-3*::GFP(mito)] - d1 2h Exercise | 53 | two-tailed Student's t vs. d1 0h | 0.2839 |
| | zcIs14[*myo-3*::GFP(mito)] - d1 4h Exercise | 56 | two-tailed Student's t vs. d1 0h | 0.0004 |
| | zcIs14[m*yo-3*::GFP(mito)] - d1 24h Recovery | 37 | two-tailed Student's t vs. d1 0h | 0.7309 |
| | | | two-tailed Student's t vs. d1 4h | 0.0010 |
| 1E (3) | zcIs14[*myo-3*::GFP(mito)] - d5 0h | 67 | | |
| | zcIs14[*myo-3*::GFP(mito)] - d5 1h Exercise | 65 | two-tailed Student's t vs. d5 0h | 0.2880 |
| | zcIs14[*myo-3*::GFP(mito)] - d5 2h Exercise | 69 | two-tailed Student's t vs. d5 0h | 0.0062 |
| | zcIs14[*myo-3*::GFP(mito)] - d5 4h Exercise | 70 | two-tailed Student's t vs. d5 0h | <.0001 |
| | zcIs14[*myo-3*::GFP(mito)] - d5 24h Recovery | 64 | two-tailed Student's t vs. d5 0h | 0.0444 |
| | | | two-tailed Student's t vs. d5 4h | 0.0002 |
| 1E (3) | zcIs14[*myo-3*::GFP(mito)] - d10 0h | 66 | | |
| | zcIs14[*myo-3*::GFP(mito)] - d10 1h Exercise | 66 | two-tailed Student's t vs. d10 0h | 0.0932 |
| | zcIs14[*myo-3*::GFP(mito)] - d10 2h Exercise | 68 | two-tailed Student's t vs. d10 0h | <.0001 |
| | zcIs14[*myo-3*::GFP(mito)] - d10 4h Exercise | 70 | two-tailed Student's t vs. d10 0h | <.0001 |
| | zcIs14[*myo-3*::GFP(mito)] - d10 24h Recovery | 59 | two-tailed Student's t vs. d10 0h | 0.0372 |
| | | | two-tailed Student's t vs. d10 4h | <.0001 |
| 1F (3) | zcIs14[*myo-3*::GFP(mito)] - d1 Recovery rate (24h - 0h) | 56 | | |
| | zcIs14[*myo-3*::GFP(mito)] - d5 Recovery rate (24h - 0h) | 67 | two-tailed Student's t vs d1 | 0.0073 |
| | zcIs14[*myo-3*::GFP(mito)] - d10 Recovery rate (24h - 0h) | 65 | two-tailed Student's t vs d1 | 0.0050 |
| 1G (3) | zcIs14[*myo-3*::GFP(mito)] - d1 0h, 1hEx, 2hEx, 4hEx, 24hRec | 5 (257) | Linear Regression | <.0001 |
| | zcIs14[*myo-3*::GFP(mito)] - d5 0h, 1hEx, 2hEx, 4hEx, 24hRec | 5 (335) | | |
| | zcIs14[*myo-3*::GFP(mito)] - d10 0h, 1hEx, 2hEx, 4hEx, 24hRec | 5 (329) | | |
| 2B (3) | N2 - d1, d5, d10, d15 | 67, 63, 54, 49 | | |
| | *drp-1(tm1108)* - d1, d5, d10, d15 | 42, 48, 46, 48 | two-tailed Student's t vs. N2 | 0.0019 |
| | *fzo-1(tm1133)* - d1, d5, d10, d15 | 42, 42, 39, 44 | two-tailed Student's t vs. N2 | 0.0015 |
| | *drp-1(tm1108);fzo-1(tm1133)* - d1, d5, d10, d15 | 69, 70, 70, 65 | two-tailed Student's t vs. N2 | 0.0036 |
| | *drp-1(tm1108);eat-3(ad426)* - d1, d5, d10, d15 | 54, 48, 61, 57 | two-tailed Student's t vs. N2 | 0.0017 |
| 2B (3) | N2 - Overall capacity (d1+d5+d10+d15) | 233 | | |
| | *drp-1(tm1108)* - Overall capacity (d1+d5+d10+d15) | 184 | two-tailed Student's t vs. N2 | <.0001 |
| | *fzo-1(tm1133)* - Overall capacity (d1+d5+d10+d15) | 167 | two-tailed Student's t vs. N2 | <.0001 |
| | *drp-1(tm1108);fzo-1(tm1133)* - Overall capacity (d1+d5+d10+d15) | 274 | two-tailed Student's t vs. N2 | <.0001 |

| | | | | |
|---|---|---|---|---|
| | *drp-1(tm1108);eat-3(ad426)* - Overall capacity (d1+d5+d10+d15) | 220 | two-tailed Student's t vs. N2 | <.0001 |
| 2C (3) | N2 - d1 0h, 1h Ex, 2h Ex, 4h Ex, 24h Rec | 56, 55, 53, 56, 37 | | |
| | *drp-1(tm1108)* - d1 0h, 1hEx, 2hEx, 4hEx, 24hRec | 56, 64, 64, 66, 62 | two-tailed Student's t vs. N2 d1 | 0.0149 |
| | *fzo-1(tm1133)* - d1 0h, 1hEx, 2hEx, 4hEx, 24hRec | 51, 61, 61, 64, 51 | two-tailed Student's t vs. N2 d1 | <.0001 |
| | *drp-1(tm1108);fzo-1(tm1133)* - d1 0h, 1hEx, 2hEx, 4hEx, 24hRec | 61, 65, 66, 69, 62 | two-tailed Student's t vs. N2 d1 | 0.0145 |
| | *drp-1(tm1108);eat-3(ad426)* - d1 0h, 1hEx, 2hEx, 4hEx, 24hRec | 64, 65, 66, 64, 58 | two-tailed Student's t vs. N2 d1 | 0.0388 |
| 2E (3) | zcIs14[*myo-3*::GFP(mito)] - Control d1 | 67 | | |
| | zcIs14[*myo-3*::GFP(mito)] - Exercise d1 | 68 | two-tailed Student's t vs. Control d1 | 0.6006 |
| | zcIs14[*myo-3*::GFP(mito)] - Control d5 | 63 | | |
| | zcIs14[*myo-3*::GFP(mito)] - Exercise d5 | 44 | two-tailed Student's t vs. Control d5 | 0.1348 |
| | zcIs14[*myo-3*::GFP(mito)] - Control d10 | 54 | | |
| | zcIs14[*myo-3*::GFP(mito)] - Exercise d10 | 65 | two-tailed Student's t vs. Control d10 | 0.0004 |
| | zcIs14[*myo-3*::GFP(mito)] - Control d15 | 49 | | |
| | zcIs14[*myo-3*::GFP(mito)] - Exercise d15 | 59 | two-tailed Student's t vs. Control d15 | <.0001 |
| 2F (3) | zcIs14[*myo-3*::GFP(mito)] - Control d5 | 173 | | |
| | zcIs14[*myo-3*::GFP(mito)] - Exercise d5 | 147 | chi-square vs. Control d5 | 0.0449 |
| | zcIs14[*myo-3*::GFP(mito)] - Control d10 | 151 | | |
| | zcIs14[*myo-3*::GFP(mito)] - Exercise d10 | 130 | chi-square vs. Control d10 | 0.0168 |
| 2G (3) | zcIs14[*myo-3*::GFP(mito)] - Control d5, d10 | 2 (117) | | |
| | zcIs14[*myo-3*::GFP(mito)] - Exercise d5, d10 | 2 (109) | | |
| 2H (3) | *drp-1(tm1108)* - Control d1 | 52 | | |
| | *drp-1(tm1108)* - Exercise d1 | 53 | two-tailed Student's t vs. Control d1 | 0.9518 |
| | *drp-1(tm1108)* - Control d5 | 48 | | |
| | *drp-1(tm1108)* - Exercise d5 | 43 | two-tailed Student's t vs. Control d5 | 0.0502 |
| | *drp-1(tm1108)* - Control d10 | 46 | | |
| | *drp-1(tm1108)* - Exercise d10 | 43 | two-tailed Student's t vs. Control d10 | 0.4028 |
| | *drp-1(tm1108)* - Control d15 | 48 | | |
| | *drp-1(tm1108)* - Exercise d15 | 48 | two-tailed Student's t vs. Control d15 | 0.4557 |
| 2H (3) | *fzo-1(tm1133)* - Control d1 | 52 | | |
| | *fzo-1(tm1133)* - Exercise d1 | 52 | two-tailed Student's t vs. Control d1 | 0.3720 |
| | *fzo-1(tm1133)* - Control d5 | 42 | | |
| | *fzo-1(tm1133)* - Exercise d5 | 43 | two-tailed Student's t vs. Control d5 | 0.3977 |
| | *fzo-1(tm1133)* - Control d10 | 39 | | |
| | *fzo-1(tm1133)* - Exercise d10 | 40 | two-tailed Student's t vs. Control d10 | 0.9599 |
| | *fzo-1(tm1133)* - Control d15 | 44 | | |
| | *fzo-1(tm1133)* - Exercise d15 | 35 | two-tailed Student's t vs. Control d15 | 0.0752 |
| 2H (3) | *drp-1(tm1108);fzo-1(tm1133)* - Control d1 | 69 | | |
| | *drp-1(tm1108);fzo-1(tm1133)* - Exercise d1 | 69 | two-tailed Student's t vs. Control d1 | 0.9146 |
| | *drp-1(tm1108);fzo-1(tm1133)* - Control d5 | 70 | | |
| | *drp-1(tm1108);fzo-1(tm1133)* - Exercise d5 | 65 | two-tailed Student's t vs. Control d5 | 0.0699 |
| | *drp-1(tm1108);fzo-1(tm1133)* - Control d10 | 70 | | |
| | *drp-1(tm1108);fzo-1(tm1133)* - Exercise d10 | 63 | two-tailed Student's t vs. Control d10 | 0.0462 |
| | *drp-1(tm1108);fzo-1(tm1133)* - Control d15 | 65 | | |

| | | | | |
|---|---|---|---|---|
| | *drp-1(tm1108);fzo-1(tm1133)* - Exercise d15 | 75 | two-tailed Student's t vs. Control d15 | 0.0600 |
| 2H (3) | *drp-1(tm1108);eat-3(ad426)* - Control d1 | 54 | | |
| | *drp-1(tm1108);eat-3(ad426)* - Exercise d1 | 55 | two-tailed Student's t vs. Control d1 | 0.5846 |
| | *drp-1(tm1108);eat-3(ad426)* - Control d5 | 48 | | |
| | *drp-1(tm1108);eat-3(ad426)* - Exercise d5 | 64 | two-tailed Student's t vs. Control d5 | 0.1322 |
| | *drp-1(tm1108);eat-3(ad426)* - Control d10 | 61 | | |
| | *drp-1(tm1108);eat-3(ad426)* - Exercise d10 | 68 | two-tailed Student's t vs. Control d10 | 0.0085 |
| | *drp-1(tm1108);eat-3(ad426)* - Control d15 | 57 | | |
| | *drp-1(tm1108);eat-3(ad426)* - Exercise d15 | 82 | two-tailed Student's t vs. Control d15 | 0.0115 |
| 2I (2) | *sid-1(qt-9);*uthIs237*;EV(RNAi)* - Control d1 | 24 | | |
| | *sid-1(qt-9);*uthIs237*;EV(RNAi)* - Exercise d1 | 24 | two-tailed Student's t vs. Control d1 | 0.6546 |
| | *sid-1(qt-9);*uthIs237*;EV(RNAi)* - Control d10 | 24 | | |
| | *sid-1(qt-9);*uthIs237*;EV(RNAi)* - Exercise d10 | 24 | two-tailed Student's t vs. Control d10 | 0.0063 |
| 2I (2) | *sid-1(qt-9)*;uthIs237;*drp-1*(RNAi) - Control d1 | 21 | | |
| | *sid-1(qt-9)*;uthIs237;*drp-1*(RNAi) - Exercise d1 | 24 | two-tailed Student's t vs. Control d1 | 0.2646 |
| | *sid-1(qt-9)*;uthIs237;*drp-1*(RNAi) - Control d10 | 20 | | |
| | *sid-1(qt-9)*;uthIs237;*drp-1*(RNAi) - Exercise d10 | 24 | two-tailed Student's t vs. Control d10 | 0.2871 |
| 2I (2) | *sid-1(qt-9);*uthIs237*;fzo-1(RNAi)* - Control d1 | 17 | | |
| | *sid-1(qt-9);*uthIs237*;fzo-1(RNAi)* - Exercise d1 | 24 | two-tailed Student's t vs. Control d1 | 0.8048 |
| | *sid-1(qt-9);*uthIs237*;fzo-1(RNAi)* - Control d10 | 24 | | |
| | *sid-1(qt-9);*uthIs237*;fzo-1(RNAi)* - Exercise d10 | 24 | two-tailed Student's t vs. Control d10 | 0.5977 |
| 2I (2) | *sid-1(qt-9)*;uthIs237;*eat-3*(RNAi) - Control d1 | 24 | | |
| | *sid-1(qt-9)*;uthIs237;*eat-3*(RNAi) - Exercise d1 | 24 | two-tailed Student's t vs. Control d1 | 0.2244 |
| | *sid-1(qt-9)*;uthIs237;*eat-3*(RNAi) - Control d10 | 24 | | |
| | *sid-1(qt-9)*;uthIs237;*eat-3*(RNAi) - Exercise d10 | 24 | two-tailed Student's t vs. Control d10 | 0.4201 |
| 3A (2) | N2 - d1 | 47 | | |
| | *isp-1(qm150)* - d1 | 43 | two-tailed Student's t vs. N2 d1 | <.0001 |
| | *nuo-6(qm200)* - d1 | 61 | two-tailed Student's t vs. N2 d1 | <.0001 |
| | *daf-2(e1370)* - d1 | 57 | two-tailed Student's t vs. N2 d1 | 0.0004 |
| | *eat-2(ad1116)* - d1 | 48 | two-tailed Student's t vs. N2 d1 | 0.0043 |
| 3A (2) | N2 - d10 | 42 | | |
| | *isp-1(qm150)* - d10 | 33 | two-tailed Student's t vs. N2 d10 | <.0001 |
| | *nuo-6(qm200)* - d10 | 33 | two-tailed Student's t vs. N2 d10 | <.0001 |
| | *daf-2(e1370)* - d10 | 34 | two-tailed Student's t vs. N2 d10 | 0.0032 |
| | *eat-2(ad1116)* - d10 | 31 | two-tailed Student's t vs. N2 d10 | 0.5281 |
| 3B (2) | N2 - d1, d5, d10, d15 | 38, 41, 42, 37 | | |
| | CA-AAK-2 - d1, d5, d10, d15 | 49, 37, 47, 50 | two-tailed Student's t vs. N2 | 0.0261 |
| 3B (2) | N2 - Overall capacity (d1+d5+d10+d15) | 158 | | |
| | CA-AAK-2 - Overall capacity (d1+d5+d10+d15) | 183 | two-tailed Student's t vs. N2 | <.0001 |
| 3C (2) | N2 - d1 0h, 1hEx, 2hEx, 4hEx, 24hRec | 38, 37, 36, 38, 37 | | |
| | CA-AAK-2 - d1 0h, 1hEx, 2hEx, 4hEx, 24hRec | 40, 39, 40, 40, 45 | two-tailed Student's t vs. N2 d1 | 0.0094 |
| 3C (2) | N2 - d5 0h, 1hEx, 2hEx, 4hEx, 24hRec | 45, 44, 47, 46, 42 | | |
| | CA-AAK-2 - d5 0h, 1hEx, 2hEx, 4hEx, 24hRec | 45, 45, 44, 45, 47 | two-tailed Student's t vs. N2 d5 | 0.0013 |
| 3C | N2 - d10 0h, 1hEx, 2hEx, 4hEx, 24hRec | 44, 41, 44, 45, 36 | | |

| | | | | |
|---|---|---|---|---|
| (2) | CA-AAK-2 - d10 0h, 1hEx, 2hEx, 4hEx, 24hRec | 44, 45, 46, 45, 43 | two-tailed Student’s t vs. N2 d10 | 0.0005 |
| 3D (2) | CA-AAK-2 - Control d1 | 49 | | |
| | CA-AAK-2 - Exercise d1 | 40 | two-tailed Student’s t vs. Control d1 | 0.7949 |
| | CA-AAK-2 - Control d5 | 37 | | |
| | CA-AAK-2 - Exercise d5 | 47 | two-tailed Student’s t vs. Control d5 | <.0001 |
| | CA-AAK-2 - Control d10 | 47 | | |
| | CA-AAK-2 - Exercise d10 | 40 | two-tailed Student’s t vs. Control d10 | <.0001 |
| | CA-AAK-2 - Control d15 | 50 | | |
| | CA-AAK-2 - Exercise d15 | 50 | two-tailed Student’s t vs. Control d15 | 0.0079 |
| 4A (2) | N2 - d1, d5, d10, d15 | 47, 41, 42, 37 | | |
| | *aak-2(gt33)* -- d1, d5, d10, d15 | 45, 52, 44, 29 | two-tailed Student’s t vs. N2 | 0.0951 |
| 4A (2) | N2 - Overall capacity (d1+d5+d10+d15) | 167 | | |
| | *aak-2(gt33)* - Overall capacity (d1+d5+d10+d15) | 170 | two-tailed Student’s t vs. N2 | 0.0010 |
| 4B (2) | N2 - d1 0h, 1hEx, 2hEx, 4hEx, 24hRec | 38, 37, 36, 38, 40 | | |
| | *aak-2(gt33)* - d1 0h, 1hEx, 2hEx, 4hEx, 24hRec | 38, 39, 39, 38, 45 | two-tailed Student’s t vs. N2 d1 | 0.0035 |
| 4C (2) | *aak-2(gt33)* - Control d1 | 45 | | |
| | *aak-2(gt33)* - Exercise d1 | 45 | two-tailed Student’s t vs. Control d1 | 0.9269 |
| | *aak-2(gt33)* - Control d5 | 52 | | |
| | *aak-2(gt33)* - Exercise d5 | 41 | two-tailed Student’s t vs. Control d5 | 0.6691 |
| | *aak-2(gt33)* - Control d10 | 44 | | |
| | *aak-2(gt33)* - Exercise d10 | 39 | two-tailed Student’s t vs. Control d10 | 0.0003 |
| | *aak-2(gt33)* - Control d15 | 29 | | |
| | *aak-2(gt33)* - Exercise d15 | 44 | two-tailed Student’s t vs. Control d15 | <.0001 |
| 4D (3) | N2 - d1, d5, d10, d15 | 62, 61, 56, 61 | | |
| | CA-AAK-2 - d1, d5, d10, d15 | 63, 84, 69, 71 | two-tailed Student’s t vs. N2 | 0.0463 |
| | CA-AAK-2;*drp-1(tm1108)* - d1, d5, d10, d15 | 70, 90, 68, 69 | two-tailed Student’s t vs. N2 | 0.2522 |
| | | | two-tailed Student’s t vs. CA-AAK-2 | 0.0486 |
| | CA-AAK-2;*fzo-1(tm1103)* - d1, d5, d10, d15 | 82, 72, 66, 65 | two-tailed Student’s t vs. N2 | 0.0204 |
| | | | two-tailed Student’s t vs. CA-AAK-2 | 0.0023 |
| | CA-AAK-2;*drp-1(tm1108);fzo-1(tm1133)* - d1, d5, d10, d15 | 57, 60, 60, 60 | two-tailed Student’s t vs. N2 | 0.0322 |
| | | | two-tailed Student’s t vs. CA-AAK-2 | <.0001 |
| 4D (3) | N2 - Overall capacity (d1+d5+d10+d15) | 240 | | |
| | CA-AAK-2 - Overall capacity (d1+d5+d10+d15) | 287 | two-tailed Student’s t vs. N2 | <.0001 |
| | CA-AAK-2;*drp-1(tm1108)* - Overall capacity (d1+d5+d10+d15) | 297 | two-tailed Student’s t vs. N2 | 0.0992 |
| | | | two-tailed Student’s t vs. CA-AAK-2 | <.0001 |
| | CA-AAK-2;*fzo-1(tm1103)* - Overall capacity (d1+d5+d10+d15) | 285 | two-tailed Student’s t vs. N2 | <.0001 |
| | | | two-tailed Student’s t vs. CA-AAK-2 | <.0001 |
| | CA-AAK-2;*drp-1(tm1108);fzo-1(tm1133)* - Overall capacity (d1+d5+d10+d15) | 237 | two-tailed Student’s t vs. N2 | <.0001 |
| | | | two-tailed Student’s t vs. CA-AAK-2 | <.0001 |
| 4E (3) | *drp-1(tm1108)* - d1, d5, d10, d15 | 56, 68, 64, 41 | | |
| | *CA-AAK-2;drp-1(tm1108)* - d1, d5, d10, d15 | 64, 62, 66, 69 | two-tailed Student’s t vs. *drp-1* | 0.1506 |
| 4E (3) | *fzo-1(tm1133)* - d1, d5, d10, d15 | 51, 57, 52, 50 | | |
| | CA-AAK-2;*fzo-1(tm1133)* - d1, d5, d10, d15 | 67, 72, 66, 65 | two-tailed Student’s t vs. f*zo-1* | 0.2512 |
| 4E | *drp-1(tm1108);fzo-1(tm1133)* - d1, d5, d10, d15 | 61, 70, 56, 65 | | |

| | | | | |
|---|---|---|---|---|
| (3) | CA-AAK-2;*drp-1(tm1108);fzo-1(tm1133)* - d1, d5, d10, d15 | 57, 60, 60, 60 | two-tailed Student's t vs. *drp-1;fzo-1* | 0.9715 |
| S1A (2-3) | N2 - d1, d5, d10, d15 | 81, 43, 68, 48 | | |
| | *drp-1* o/e - d1, d5, d10, d15 | 39, 39, 40, 29 | two-tailed Student's t vs. N2 | 0.0075 |
| | *fzo-1* o/e - d1, d5, d10, d15 | 64, 61, 66, 43 | two-tailed Student's t vs. N2 | 0.0142 |
| S1A (2-3) | N2 - Overall capacity (d1+d5+d10+d15) | 240 | | |
| | *drp-1* o/e - Overall capacity (d1+d5+d10+d15) | 147 | two-tailed Student's t vs. N2 | <.0001 |
| | *fzo-1* o/e - Overall capacity (d1+d5+d10+d15) | 234 | two-tailed Student's t vs. N2 | <.0001 |
| S1B (3) | N2 - Fitness capacity d1 (0h+ 1hEx+ 2hEx+ 4hEx) | 220 | | |
| | *drp-1(tm1108)* - Fitness capacity d1 (0h+ 1hEx+ 2hEx+ 4hEx) | 250 | two-tailed Student's t vs. N2 d1 | <.0001 |
| | *fzo-1(tm1133)* - Fitness capacity d1 (0h+ 1hEx+ 2hEx+ 4hEx) | 237 | two-tailed Student's t vs. N2 d1 | <.0001 |
| | *drp-1(tm1108);fzo-1(tm1133)* - Fitness capacity d1 (0h+ 1hEx+ 2hEx+ 4hEx) | 261 | two-tailed Student's t vs. N2 d1 | <.0001 |
| | *drp-1(tm1108);eat-3(ad426)* - Fitness capacity d1 (0h+ 1hEx+ 2hEx+ 4hEx) | 259 | two-tailed Student's t vs. N2 d1 | 0.0027 |
| S1C (3) | N2 - d1 Recovery rate (24h - 0h) | 56 | | |
| | *drp-1(tm1108)* - d1 Recovery rate (24h - 0h) | 56 | two-tailed Student's t vs. N2 d1 | 0.0046 |
| | *fzo-1(tm1133)* - d1 Recovery rate (24h - 0h) | 52 | two-tailed Student's t vs. N2 d1 | 0.0470 |
| | *drp-1(tm1108);fzo-1(tm1133)* - d1 Recovery rate (24h - 0h) | 61 | two-tailed Student's t vs. N2 d1 | 0.9375 |
| | *drp-1(tm1108);eat-3(ad426)* - d1 Recovery rate (24h - 0h) | 65 | two-tailed Student's t vs. N2 d1 | 0.1933 |
| S2A (2) | N2 - d1 0h, 1hEx, 2hEx, 4hEx, 24hRec | 37, 50, 50, 51, 42 | | |
| | *isp-1(qm150)* - d1 0h, 1hEx, 2hEx, 4hEx, 24hRec | 48, 47, 49, 49, 46 | two-tailed Student's t vs. N2 d1 | 0.0013 |
| | *nuo-6(qm200)* - d1 0h, 1hEx, 2hEx, 4hEx, 24hRec | 44, 44, 43, 45, 51 | two-tailed Student's t vs. N2 d1 | 0.0036 |
| S2A (2) | N2 - Fitness capacity d1 (0h+ 1hEx+ 2hEx+ 4hEx) | 188 | | |
| | *isp-1(qm150)* - Fitness capacity d1 (0h+ 1hEx+ 2hEx+ 4hEx) | 193 | two-tailed Student's t vs. N2 d1 | <.0001 |
| | *nuo-6(qm200)* - Fitness capacity d1 (0h+ 1hEx+ 2hEx+ 4hEx) | 176 | two-tailed Student's t vs. N2 d1 | <.0001 |
| S2B (2) | N2 - d1 0h, 1hEx, 2hEx, 4hEx, 24hRec | 37, 50, 50, 51, 42 | | |
| | *daf-2(e1370)* - d1 0h, 1hEx, 2hEx, 4hEx, 24hRec | 47, 48, 47, 49, 45 | two-tailed Student's t vs. N2 d1 | 0.0080 |
| | *eat-2(ad1116)* - d1 0h, 1hEx, 2hEx, 4hEx, 24hRec | 34, 41, 36, 40, 33 | two-tailed Student's t vs. N2 d1 | 0.0193 |
| S2B (2) | N2 - Fitness capacity d1 (0h+ 1hEx+ 2hEx+ 4hEx) | 188 | | |
| | *daf-2(e1370)* - Fitness capacity d1 (0h+ 1hEx+ 2hEx+ 4hEx) | 191 | two-tailed Student's t vs. N2 d1 | <.0001 |
| | *eat-2(ad1116)* - Fitness capacity d1 (0h+ 1hEx+ 2hEx+ 4hEx) | 151 | two-tailed Student's t vs. N2 d1 | 0.0001 |
| S2C (2) | N2 - d1 Recovery rate (24h - 0h) | 37 | | |
| | *isp-1(qm150)* - d1 Recovery rate (24h - 0h) | 48 | two-tailed Student's t vs. N2 d1 | 0.2348 |
| | *nuo-6(qm200)* - d1 Recovery rate (24h - 0h) | 44 | two-tailed Student's t vs. N2 d1 | <.0001 |
| S2D (2) | N2 - d1 Recovery rate (24h - 0h) | 37 | | |
| | *daf-2(e1370)* - d1 Recovery rate (24h - 0h) | 47 | two-tailed Student's t vs. N2 d1 | 0.0002 |
| | *eat-2(ad1116)* - d1 Recovery rate (24h - 0h) | 34 | two-tailed Student's t vs. N2 d1 | 0.3982 |
| S2E (2) | *isp-1(qm150)* - Control d1 | 21 | | |
| | *isp-1(qm150)* - Exercise d1 | 22 | two-tailed Student's t vs. Control d1 | 0.9172 |
| | *isp-1(qm150)* - Control d10 | 33 | | |
| | *isp-1(qm150)* - Exercise d10 | 33 | two-tailed Student's t vs. Control d10 | 0.5897 |
| S2E (2) | *nuo-6(qm200)* - Control d1 | 31 | | |
| | *nuo-6(qm200)* - Exercise d1 | 32 | two-tailed Student's t vs. Control d1 | 0.6668 |
| | *nuo-6(qm200)* - Control d10 | 33 | | |
| | *nuo-6(qm200)* - Exercise d10 | 19 | two-tailed Student's t vs. Control d10 | 0.4912 |

| | | | | |
|---|---|---|---|---|
| S2F (2) | *daf-2(e1370)* - Control d1 | 29 | | |
| | *daf-2(e1370)* - Exercise d1 | 28 | two-tailed Student's t vs. Control d1 | 0.6613 |
| | *daf-2(e1370)* - Control d10 | 34 | | |
| | *daf-2(e1370)* - Exercise d10 | 24 | two-tailed Student's t vs. Control d10 | 0.5942 |
| S2F (2) | *eat-2(ad1116)* - Control d1 | 29 | | |
| | *eat-2(ad1116)* - Exercise d1 | 28 | two-tailed Student's t vs. Control d1 | 0.5150 |
| | *eat-2(ad1116)* - Control d10 | 31 | | |
| | *eat-2(ad1116)* - Exercise d10 | 23 | two-tailed Student's t vs. Control d10 | 0.4225 |
| S2G (2) | N2 - Fitness capacity d1 (0h+ 1hEx+ 2hEx+ 4hEx) | 149 | | |
| | CA-AAK-2 - Fitness capacity d1 (0h+ 1hEx+ 2hEx+ 4hEx) | 159 | two-tailed Student's t vs. N2 d1 | <.0001 |
| S2G (2) | N2 - Fitness capacity d5 (0h+ 1hEx+ 2hEx+ 4hEx) | 182 | | |
| | CA-AAK-2 - Fitness capacity d5 (0h+ 1hEx+ 2hEx+ 4hEx) | 179 | two-tailed Student's t vs. N2 d5 | <.0001 |
| S2G (2) | N2 - Fitness capacity d10 (0h+ 1hEx+ 2hEx+ 4hEx) | 174 | | |
| | CA-AAK-2 - Fitness capacity d10 (0h+ 1hEx+ 2hEx+ 4hEx) | 180 | two-tailed Student's t vs. N2 d10 | <.0001 |
| S2H (2) | N2 - d1 Recovery rate (24h - 0h) | 38 | | |
| | CA-AAK-2 - d1 Recovery rate (24h - 0h) | 40 | two-tailed Student's t vs. N2 d1 | 0.1443 |
| S2H (2) | N2 - d5 Recovery rate (24h - 0h) | 45 | | |
| | CA-AAK-2 - d5 Recovery rate (24h - 0h) | 45 | two-tailed Student's t vs. N2 d5 | 0.0264 |
| S2H (2) | N2 - d10 Recovery rate (24h - 0h) | 44 | | |
| | CA-AAK-2 - d10 Recovery rate (24h - 0h) | 44 | two-tailed Student's t vs. N2 d10 | 0.0155 |
| S3A (2) | N2 - Fitness capacity d1 (0h+ 1hEx+ 2hEx+ 4hEx) | 149 | | |
| | *aak-2(gt33)* - Fitness capacity d1 (0h+ 1hEx+ 2hEx+ 4hEx) | 154 | two-tailed Student's t vs. N2 d1 | <.0001 |
| S3B (2) | N2 - d1 Recovery rate (24h - 0h) | 38 | | |
| | *aak-2(gt33)* - d1 Recovery rate (24h - 0h) | 38 | two-tailed Student's t vs. N2 d1 | 0.0010 |
| S3C (3) | *drp-1(tm1108)* - Overall capacity (d1+d5+d10+d15) | 229 | | |
| | *CA-AAK-2;drp-1(tm1108)* - Overall capacity (d1+d5+d10+d15) | 261 | two-tailed Student's t vs. *drp-1* | 0.5522 |
| S3C (3) | *fzo-1(tm1133)* - Overall capacity (d1+d5+d10+d15) | 210 | | |
| | *CA-AAK-2;fzo-1(tm1133)* - Overall capacity (d1+d5+d10+d15) | 270 | two-tailed Student's t vs. *fzo-1* | <.0001 |
| S3C (3) | *drp-1(tm1108);fzo-1(tm1133)* - Overall capacity (d1+d5+d10+d15) | 252 | | |
| | *CA-AAK-2;drp-1(tm1108);fzo-1(tm1133)* - Overall capacity (d1+d5+d10+d15) | 237 | two-tailed Student's t vs. *drp-1;fzo-1* | 0.7733 |
